# Geometry-dependent mRNA delivery with programmable RNA-DNA nanostructures

**DOI:** 10.64898/2026.09.19.752892

**Authors:** Mengxi Zheng, Abhisek Dwivedy, Dhanush Gandavadi, Wei Hong, Hyeongjun Cho, Natalie Shkolnik, Xing Wang

## Abstract

Efficient intracellular delivery of neoantigen-encoded mRNAs is central to the development of effective cancer vaccines and immunotherapies. However, how the nanoscale geometry of delivery carriers impacts mRNA delivery remains poorly understood, largely because conventional delivery platforms do not readily permit systematic variation of carrier architecture while maintaining comparable chemical composition. Here, we develop a modular RNA-DNA hybrid nanoplatform that enables controlled investigation of geometry-dependent mRNA delivery. Guided by a unified design principle, we construct a series of two- and three-dimensional nanostructures with programmable architectures and systematically compare their delivery performance using consistent functional readouts. We further investigate how mRNA embedding strategies and aptamer-mediated functionalization influence intracellular delivery efficiency. By providing precise control over carrier geometry, this platform establishes a framework for elucidating structure-function relationships in mRNA delivery and offers a versatile strategy for engineering targeted nucleic acid delivery systems.

## INTRODUCTION

Lipid nanoparticles (LNPs) have demonstrated the transformative impact of mRNA delivery, most notably through the clinical success of LNP-formulated mRNA vaccines against COVID19^1–4^. Beyond infectious diseases, efficient and targeted intracellular delivery of mRNA is essential for emerging applications in the development of cancer vaccination and immunotherapy. Accordingly, extensive efforts have focused on optimizing LNP composition, particularly ionizable lipids, surface properties, and formulation parameters to improve mRNA stability, cellular uptake, intracellular trafficking, and cytosolic release^2, 5–7^. In comparison, the role of carrier geometry in determining the fate and functional delivery of mRNA remains considerably less understood.

Recent studies have suggested that the geometry of a nanoscale carrier could potentially influence its payload delivery efficiency and intracellular processing^8–11^. However, establishing structure-function relationships has been challenging because changes in conventional carrier geometry, such as LNP, are often accompanied by changes in size, surface chemistry, composition, or other physicochemical properties. A more systematic comparison therefore requires a platform in which geometry can be independently programmed while other design parameters are maintained as consistently as possible. Such a capability would provide an important foundation for understanding how nanoscale architecture of a carrier regulates intracellular mRNA delivery.

DNA nanotechnology provides a powerful framework for addressing this challenge. Through programmable base pairing, DNA can be assembled into well-defined structures with precise control over nanoscale dimensions, curvatures, and spatial organization^12–14^. They have demonstrated great stability in buffers, cell culture media, cell lysates, and blood^15–24^, which can be further enhanced or finely tuned through surface coating^18, 25–27^, UV crosslinking^24^, or self-assembly in Mg^2+^-free solutions^28^ or in aqueous ionic liquid solutions^29^. Importantly, distinct geometries can be generated from related molecular components, providing an opportunity to examine geometric effects while avoiding variations in chemical composition. DNA nanostructures have been explored as carriers for nucleic acids and other therapeutic cargos^30, 31^. However, a systematic evaluation of how defined carrier geometries affect mRNA delivery within a unified molecular framework remains largely unexplored.

In this work, we develop a modular RNA-DNA hybrid nanoplatform to systematically investigate geometry-dependent mRNA delivery (**Fig. 1**). Using a common design principle, we construct a series of two- and three-dimensional nanostructures that differ in architecture while retaining a closely related molecular composition. This design enables direct comparison of their cellular uptake and functional mRNA delivery using consistent experimental readouts. Beyond geometric control, the platform allows the mRNA to be incorporated through programmable embedding strategies and provides a defined docking site for aptamer integration, enabling further modulation of cellular targeting and delivery. Notably, the nanostructures can be assembled from a small number of short nucleic acid oligonucleotides, providing a relatively simple and economical approach that is particularly well suited for short mRNA cargos, including neoantigen-encoding sequences. Together, this platform provides a controlled experimental framework for dissecting geometry-dependent mRNA delivery and establishes design principles for engineering programmable nucleic acid delivery systems.

**Figure 1.**
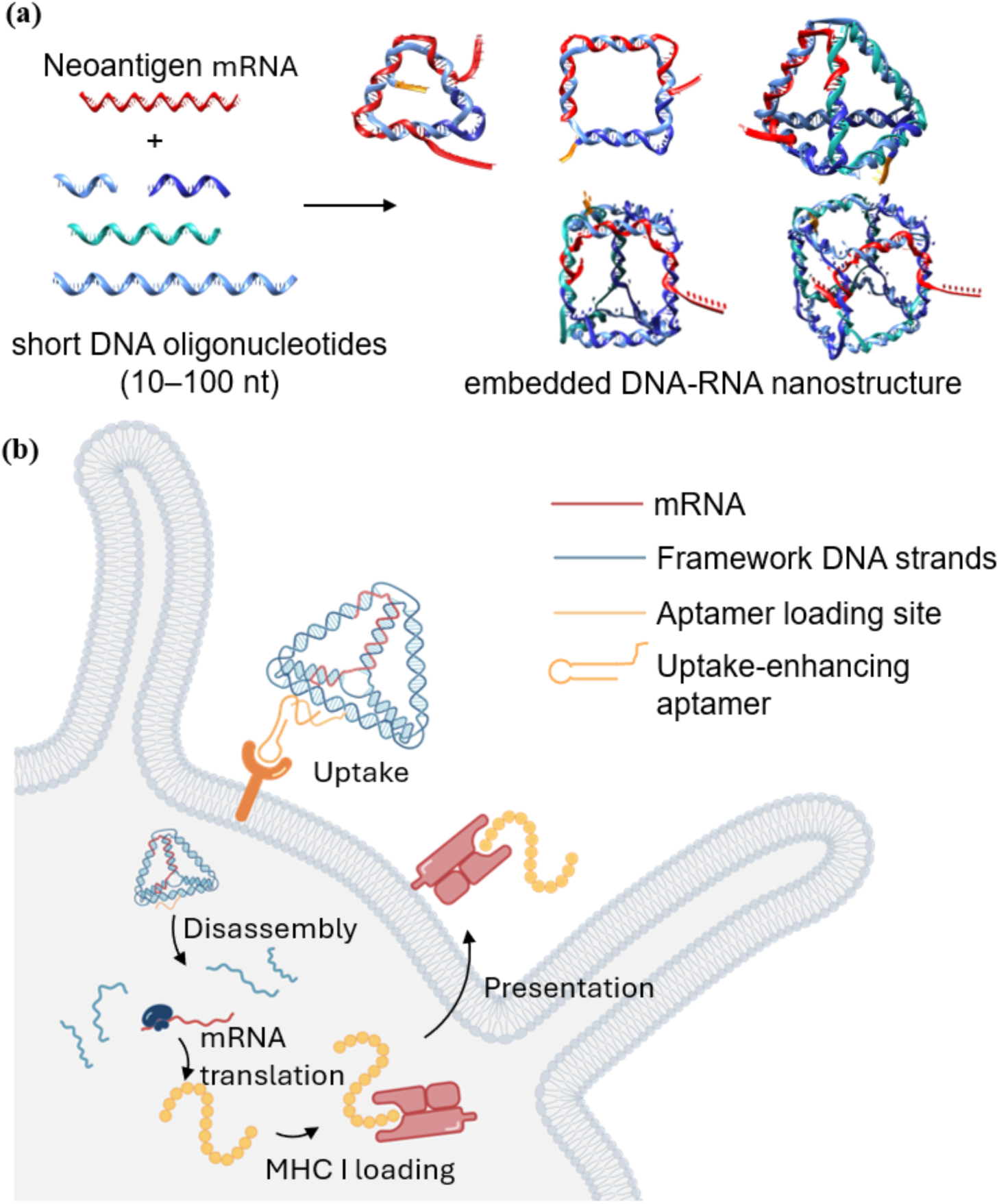
Schematic illustration of the modular and programmable mRNA-DNA nanoplatform for mRNA delivery. **(a)** Assembly of neoantigen mRNA with short DNA oligonucleotides (10 - 100 nt) into an embedded RNA-DNA nanostructure. **(b)** Cellular uptake, disassembly, and subsequent mRNA translation leading to antigen presentation via the MHC I pathway.

## RESULTS

### Design Principles

To demonstrate our platform, we selected ovalbumin mRNA (encoding the OVA Peptide 1, amino acid sequence-SIINFEKL) as the model antigen, as it is widely used in immunological studies with established assays for evaluating the efficiency of mRNA delivery and expression.

By testing vaccines against established “model” tumors like B16-OVA (B-16 melanoma line expression OVA peptide 1), scientists can reliably validate new delivery platforms against decades of existing benchmark data. In addition, neoantigens are typically encoded by short sequences of similar size to OVA mRNA^32–34^. The successful design of a compact RNA-DNA nanoplatform for OVA mRNA can be readily extended to neoantigen encoding mRNA sequences for future immunological studies. RNA sequences are prone to rapid degradation in physiological conditions. Many chemical modifications of RNA reported in the past have been utilized towards improving RNA stability and nuclease resistance. However, most modifications require expertise in click chemistry with low synthetic yields. As an alternative, we explored viral RNA motifs known as xrRNA that confer resistance to exoribonucleases^35–38^. We included xrRNA like motifs into the design of the OVA mRNA to confer enhanced exoribonuclease resistance. While OVA and neoantigen mRNA generated using this design have been reported in our previous work, the design principles are reported here^39, 40^.

In our design, the main body of OVA mRNA is embedded within the nanostructure framework to form an mRNA-DNA hybrid. The DNA-based nanostructure backbones provide a much more stable and less expensive framework than pure RNA made nanostructures. Each mRNA-DNA interaction domain is designed to be at least 8-nucleotides (nt) in length to ensure the thermostability during delivery. At the vertices of each nanostructure, 1-5 nt of OVA mRNA are left as free spacers or loops, depending on the local geometry, to allow bending and provide flexibility, thereby avoiding over-constraining given the uncertain structural details of RNA-DNA hybridizations. Although most OVA mRNA is embedded as RNA-DNA hybrids, the first 6 nucleotides of the mRNA are always left unpaired to ensure that translation is not inhibited. If the 5′ end is involved in complex secondary structures, RNA translation is often inhibited. Additionally, the 5’ UTR may form xrRNA like motifs, thus conferring protection from ribonucleases.

From a design perspective, this nanoplatform offers several advantages. Importantly, because it follows a fixed designing workflow, it can be readily adapted to actual neoantigen mRNAs. Secondly, this framework-shaped architecture provides structural diversity, as different geometries can be generated by rearranging edges and vertices, enabling the optimization of delivery efficiency originating from shape-dependent factors. Finally, the nanostructure edges are sufficiently long (20-30 base pairs) that nick points can be introduced flexibly, allowing each construct to be assembled from a single mRNA and only a few DNA oligonucleotides, which makes the nanoplatform cost-effective.

### Structural Characterization

To exploit the programmability of the hybrid RNA-DNA nanostructures, we designed 5 distinct OVA mRNA delivery carriers with different geometries: two 2D structures (triangle and square) and three 3D structures (tetrahedron, triangular prism, and cube). All nanostructures were constructed following the same design principles. Each carrier consists of a single OVA mRNA strand together with a few DNA oligonucleotides. More specifically, the triangle and square included two DNA oligonucleotides in addition to the OVA mRNA, the tetrahedron was formed from one OVA mRNA and five DNA strands, the triangular prism from six DNA strands, and the cube from eight DNA strands. The formation of each structure is confirmed by both polyacrylamide gel electrophoresis (PAGE) and atomic force microscope (AFM) imaging (**Fig. 2** and **Fig. S1-4**). The enlarged views highlight the defined geometric features of each nanostructure, confirming successful formation at the designed dimensions.

**Figure 2.**
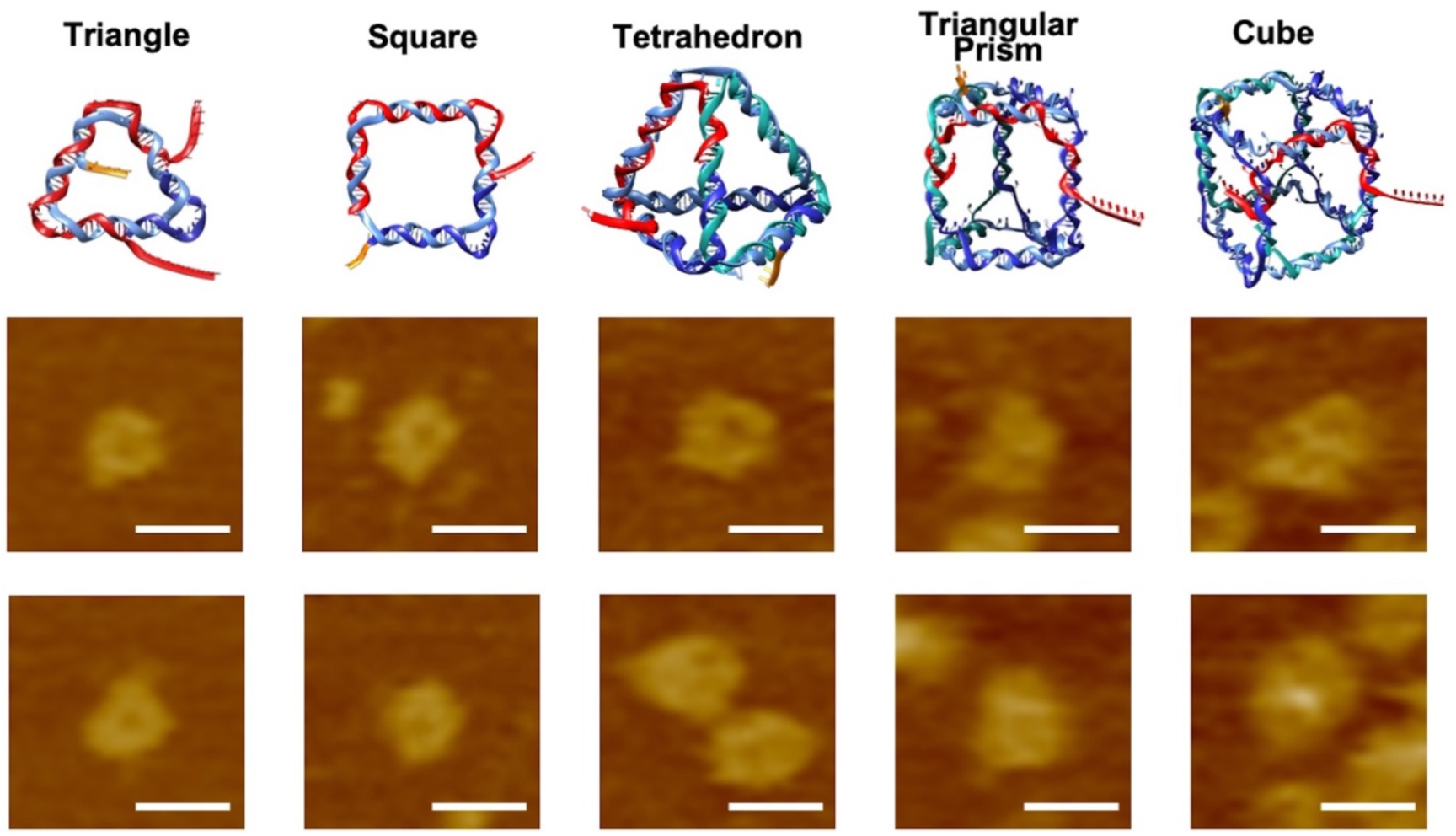
AFM Characterization of mRNA-DNA nanostructures with different geometries. Representative zoom-in AFM images of five RNA-DNA nanostructures, triangle, square, tetrahedron, triangular prism, and cube, assembled from OVA mRNA and short DNA oligonucleotides (10-100 nt). (Scale Bar: 20 nm)

Having confirmed the successful assembly of these nanostructures, we evaluated the performance of these nanostructures in mRNA delivery and expression efficiency in dendritic cells. To assess the delivery and translation efficiency of OVA mRNA, dendritic cells (DCs) were transfected with the mRNA-DNA nanostructures, and the surface presentation of OVA-derived peptide on MHC I molecules was quantified by flow cytometry (**Fig. 3a**). To ensure that geometry was evaluated as an independent variable, we established a unified and optimized delivery configuration prior to systematic comparison. Specifically, we examined different mRNA loading architectures within the nanostructure framework as well as varying mRNA dosages to identify conditions that maximize functional readout. In addition, a programmable aptamer docking site was incorporated at predefined positions in each construct to enable consistent and site-specific functionalization, thereby enhancing delivery performance while preserving the underlying geometric framework.

**Figure 3.**
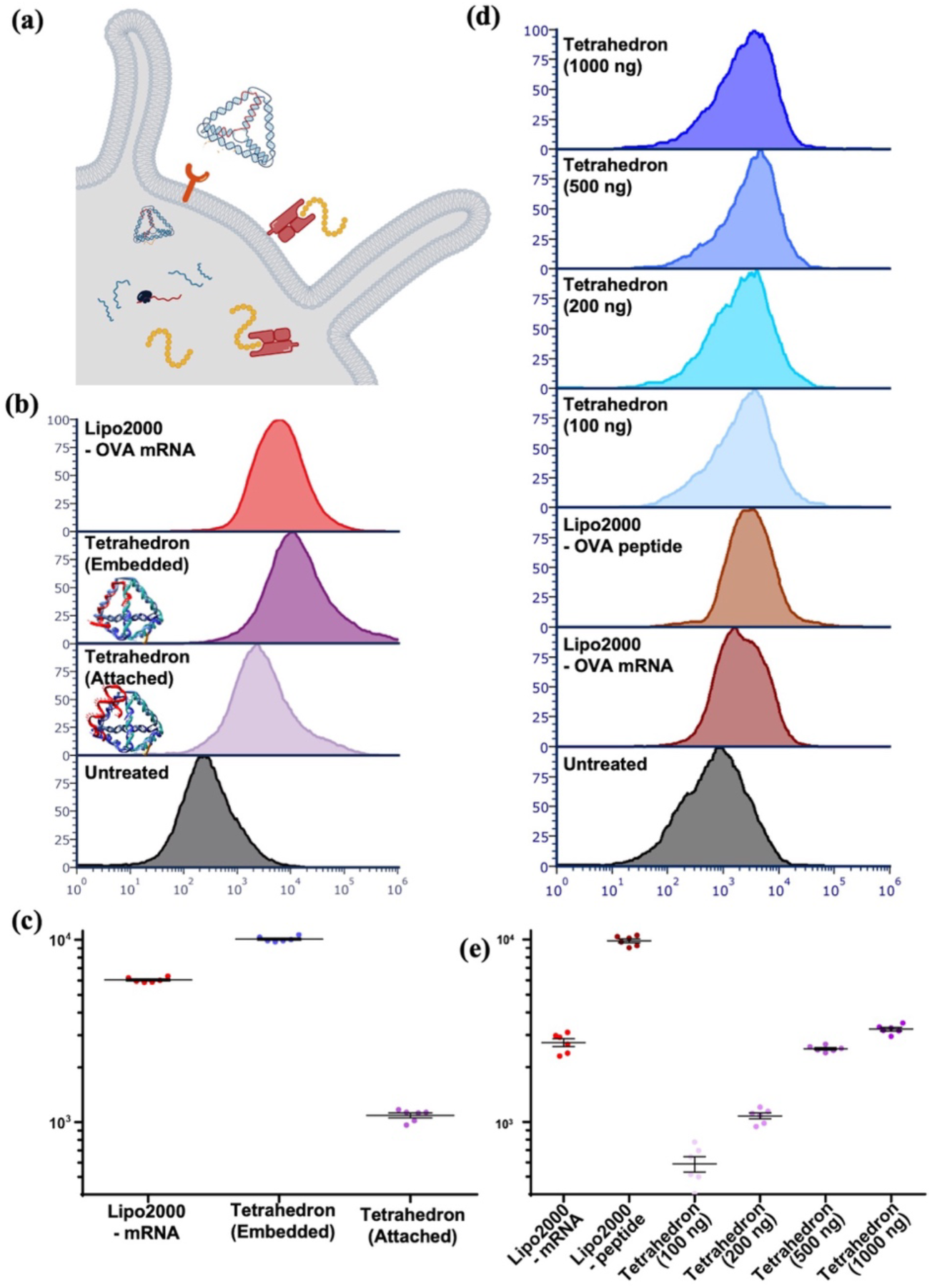
Functional evaluation of the mRNA-DNA nanoplatform without aptamer. (**a**) Experimental workflow for characterizing mRNA delivery and expression in dendritic cells without aptamer. (**b**) Representative flow cytometry histograms comparing embedded and attached mRNA configurations in tetrahedral carriers. (**c**) Quantification of background-subtracted mean fluorescence intensity (MFI) from independent replicates comparing embedded and attached configurations. (**d**) Representative flow cytometry histograms showing mRNA dose optimization using the embedded tetrahedral construct. (**e**) Corresponding quantitative analysis of background-subtracted MFI values identifying 500 ng as the optimal dosage.

### Validation of Embedded mRNA Architecture and Does Optimization

We first compared two distinct mRNA loading architectures within the tetrahedral framework (**Fig. 3b-c**). In the embedded configuration, the mRNA strand was structurally integrated into the DNA nanostructure and served as an integral component of the framework. In this design, the cargo itself constitutes part of the carrier architecture, and therefore requires geometry-specific sequence design for each construct. In contrast, in a so-called attached configuration, the mRNA was tethered to the nanostructure through several complementary base pairs at its 5′ terminus. This strategy is more modular and universally applicable, as the same mRNA strand can in principle be attached to different carrier geometries without redesigning the structural framework.

Under identical transfection conditions, the embedded architecture resulted in significantly higher levels of antigen presentation compared to the surface-attached counterpart (**Fig. 3b-c**), indicating that spatial integration of mRNA within the nanostructure enhances functional delivery efficiency. We speculate that partial hybridization between the mRNA and DNA strands within the embedded framework may provide structural stabilization and confer a better resistance against nuclease-mediated degradation. Based on these findings, the embedded design was adopted for all subsequent geometry comparisons.

We next optimized the mRNA dosage using the embedded tetrahedral construct as a representative carrier. Increasing amounts of OVA mRNA were tested, and antigen presentation was quantified by flow cytometry. Delivery efficiency increased with dose up to 500 ng and plateaued thereafter (**Fig. 3d-e**), establishing 500 ng as the standardized dosage for subsequent experiments.

### Aptamer Integration

To further enhance delivery performance while preserving geometric control, a programmable aptamer docking site was incorporated at predefined positions in each nanostructure. An in house generated CD33 targeting aptamer was employed for site-specific functionalization. CD33 is a sialic acid-binding lectin (Siglec) primarily expressed on myeloid cells, including dendritic cells^41^. This also enables modular integration of targeting ligands without altering the underlying structural framework (**Fig. 4a**).

**Figure 4.**
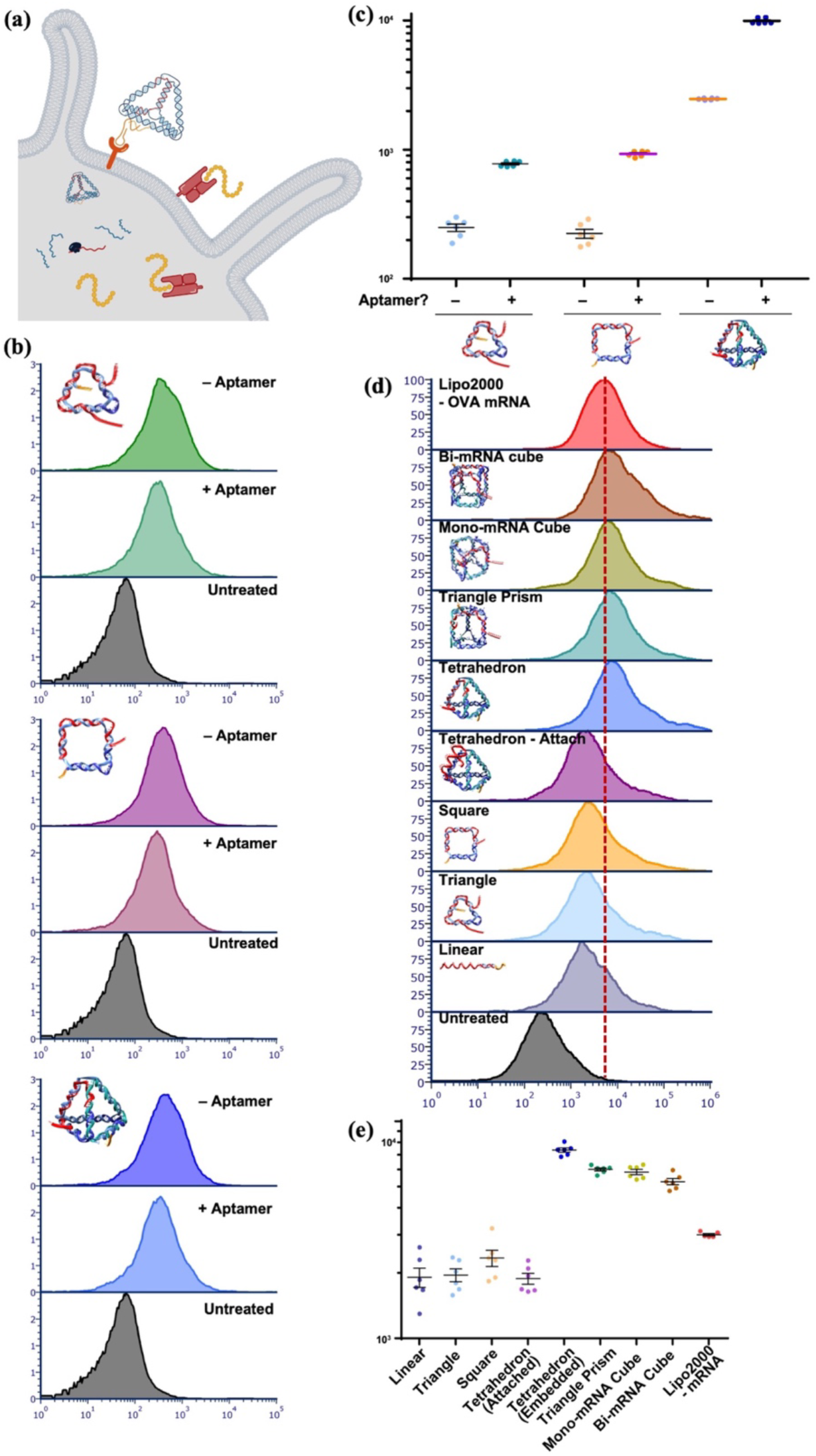
Aptamer- and geometry-dependent mRNA-DNA nanoplatform delivery. (**a**) Experimental workflow for characterizing mRNA delivery and expression in dendritic cells with aptamer. (**b**) Representative histograms comparing nanostructures with and without CD33 aptamer functionalization. (**c**) Corresponding quantitative analysis of background-subtracted MFI values demonstrating enhanced delivery upon aptamer integration. (**d**) Representative histograms showing geometry-dependent mRNA delivery efficiency across linear, two-dimensional (triangle and square), and three-dimensional (tetrahedron, triangular prism, mono-mRNA cube, and bi-mRNA cube) nanostructures under unified conditions. (**e**) Corresponding quantitative analysis of background-subtracted MFI values highlighting the hierarchical trend of 3D > 2D > linear structures.

To evaluate the functional contribution of aptamer incorporation, we compared constructs with and without the CD33 aptamer under otherwise identical conditions. The aptamer-functionalized nanostructures exhibited comparatively enhanced antigen presentation relative to their non-functionalized counterparts, demonstrating that programmable ligand integration can further improve delivery efficiency (**Fig. 4b-c**). These results confirm the modularity of the RNA-DNA hybrid platform and its compatibility with targeted functional enhancement. Unless otherwise specified, all subsequent geometry-dependent delivery studies were conducted using embedded, aptamer-functionalized nanostructures at the optimized mRNA dosage.

### Shape-Dependent Delivery Efficiency

Having established a unified delivery configuration, we next systematically compared the mRNA delivery performance of nanostructures with distinct geometries. Despite sharing identical chemical composition, embedded loading architecture, aptamer functionalization, and mRNA dosage, the constructs exhibited clear geometry-dependent differences in antigen presentation efficiency (**Fig. 4d-e**). Overall, three-dimensional (3D) nanostructures consistently outperformed two-dimensional (2D) constructs. Among the tested geometries, compact 3D architectures such as the tetrahedron, triangular prism, and cube variants demonstrated markedly higher antigen presentation levels compared to planar triangle and square structures. In contrast, the linear configuration exhibited substantially reduced delivery efficiency, approaching that of untreated controls.

Among the three-dimensional constructs, two cube variants were evaluated that differed in mRNA valency: a mono-mRNA cube containing a single embedded mRNA strand and a bi-mRNA cube incorporating two mRNA strands within the same geometric framework. This design enabled assessment of whether increasing cargo valency within an identical structural scaffold enhances delivery efficiency. Notably, the mono- and bi-mRNA cube constructs exhibited comparable antigen presentation levels, indicating that geometry rather than cargo multiplicity primarily governs functional performance under these conditions. Importantly, because the total mRNA dosage was held constant, the bi-mRNA cube achieves equivalent delivery output while potentially reducing the number of RNA-DNA hybrid nanostructures required per unit of mRNA, thereby minimizing the introduction of exogenous DNA framework material.

This hierarchical trend, 3D > 2D > linear, suggests that increasing structural dimensionality and spatial confinement enhance functional mRNA delivery. Notably, the embedded tetrahedral construct achieved one of the highest expression levels while requiring a relatively small number of DNA strands, indicating that compact and structurally balanced geometries may provide an optimal spatial environment for intracellular processing. Together, these results demonstrate that carrier geometry, beyond cargo loading alone, is a key parameter regulating mRNA delivery efficiency.

We also considered whether the different delivery efficiencies could be attributed to differences in structural stability. Melting analysis showed that the linear mRNA-DNA hybrid had a melting temperature (Tm) closer to the physiological temperature, whereas the 2D triangle and square exhibited much higher Tm values of approximately 75 °C (**Figure S6**). The 3D tetrahedron and cube showed intermediate Tm values of approximately 55-60 °C. In parallel, the length of continuous mRNA-DNA hybridization also varied among the structures, with 18 bp in the linear construct, 54-60 bp in the 2D structures, and 11-22 bp in the 3D structures. However, neither the Tm values nor the continuous hybridization lengths correlated with the observed delivery efficiency. In particular, the highly stable 2D structures did not show the highest antigen presenting performance, whereas the 3D structures exhibited stronger delivery despite their intermediate thermal stability and shorter continuous hybridization domains. These results suggest that structural stability alone is not the dominant determinant of delivery efficiency. Instead, the improved performance of the 3D constructs is more consistent with a geometry-associated effect, likely involving nanoscale topology, compactness, and spatial organization of the mRNA-DNA hybrid assembly. Thus, the melting data serve as an important control showing that the enhanced delivery of 3D mRNA-DNA hybrids cannot be explained simply by higher duplex stability or longer mRNA-DNA hybridization. Rather, geometry-associated structural features appear to play a major role in regulating biological delivery.

### Generalization of Geometry Effects to Longer mRNA Cargo

To further examine whether the observed geometry-dependent delivery behavior is conserved across different experimental systems, a longer mRNA encoding small ultra-red fluorescent protein (smURFP) was employed to construct mRNA-DNA nanostructures using a previously reported RNA-DNA hybrid origami strategy^42^, in which the RNA served as the scaffold strand and multiple short DNA staples folded it into a defined framework (**Fig. 5a**). In addition to the 3D tetrahedral architecture, a 2D rectangular nanostructure was also designed using the same smURFP mRNA scaffold with an alternative set of DNA staples, enabling direct comparison of geometric effects while maintaining identical mRNA composition. In both constructs, an aptamer docking site was incorporated at selected vertices to allow optional targeting functionalization.

**Figure 5.**
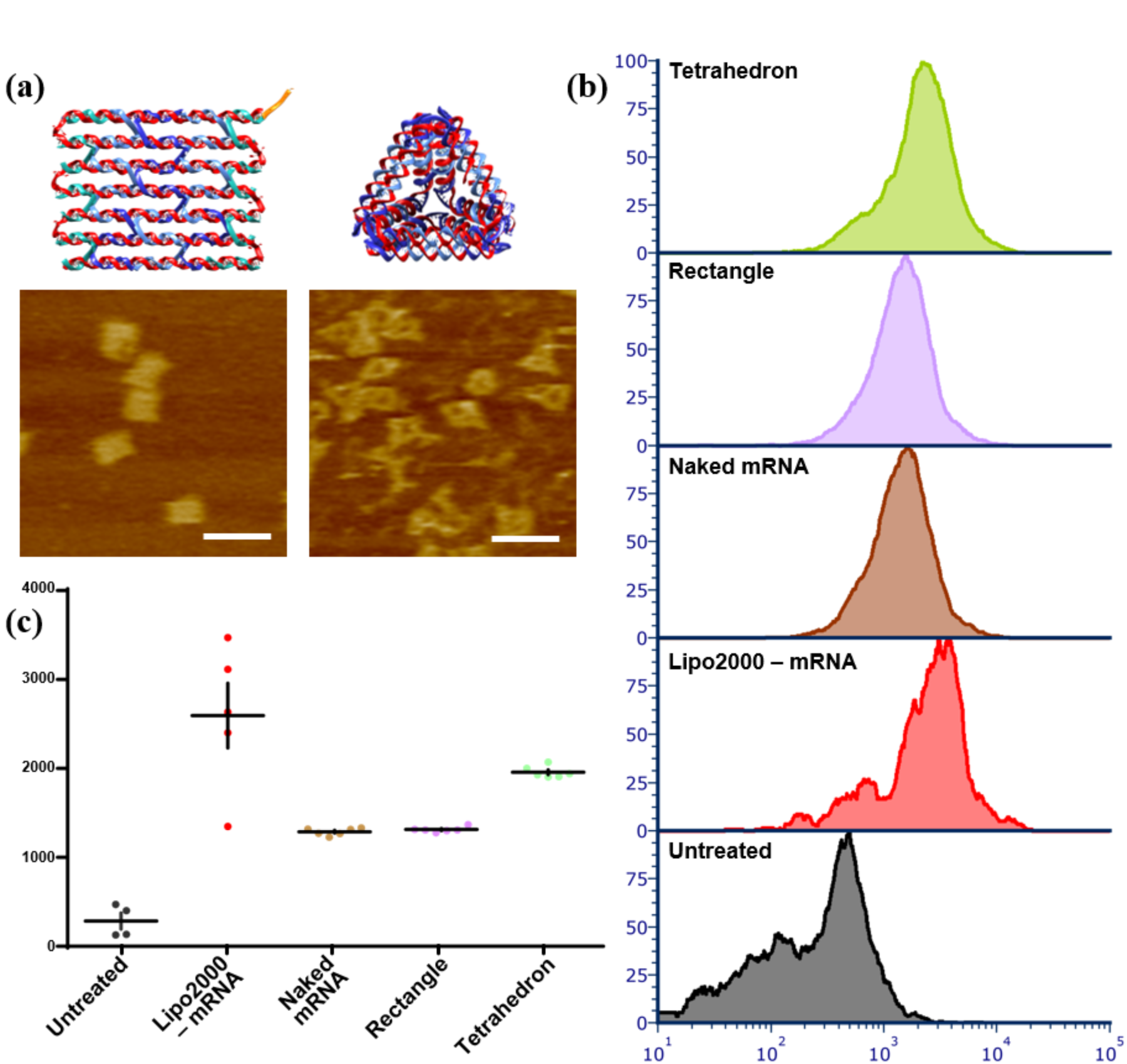
Extension of the mRNA-DNA nanoplatform to longer mRNA cargo. (**a**) Schematic illustration of the mRNA-DNA hybrid origami design using smURFP mRNA as the scaffold strand folded by short DNA staples into defined nanostructures. (**b**) Representative flow cytometry histograms showing smURFP expression in dendritic cells following transfection with naked mRNA, tetrahedral, and rectangular mRNA-DNA nanostructures. (**c**) Quantitative analysis of background-subtracted mean fluorescence intensity (MFI) from independent replicates, comparing geometry-dependent delivery efficiency of smURFP mRNA constructs.

Despite the substantially increased mRNA length relative to OVA, both tetrahedral and rectangular mRNA-DNA nanostructures assembled successfully, as confirmed by PAGE and AFM imaging (**Fig. 5a**). Upon transfection into THP-1 cells, both structured constructs exhibited markedly enhanced fluorescence expression compared to naked smURFP mRNA, indicating that structural organization of the mRNA has improved functional delivery efficiency (**Fig. 5b**). Notably, the tetrahedral architecture produced comparatively higher expression levels than the rectangular structure, consistent with the geometry-dependent trend observed in the OVA system (**Fig. 5c**). These results demonstrate that the RNA-DNA hybrid platform is compatible with longer mRNA scaffolds and that the influence of carrier geometry on delivery efficiency persists across different mRNA cargos and experimental settings. Together, these findings support a model in which nanostructure geometry plays one of the central roles in governing functional mRNA delivery.

## DISCUSSION

The influence of nanoparticle geometry on the delivery of small-molecule drugs, mRNA, and other biomacromolecules has been increasingly recognized^43–45^. However, establishing structure-function relationships remains challenging because, in conventional delivery systems, changes in particle geometry are often accompanied by variations in chemical composition, surface properties, and formulation parameters. Moreover, the range of geometries that can be readily generated and compared within a single platform is often limited. These constraints have made it difficult to systematically isolate the contribution of carrier geometry to intracellular delivery.

Here, we exploit the programmability of nucleic acid nanotechnology to construct a series of well-defined nanostructures with distinct two- and three-dimensional geometries while maintaining closely related chemical composition and surface characteristics^11, 46, 47^. This design enables parallel comparison of their delivery performance within a unified experimental framework. Our results reveal a clear geometry-dependent trend, with 3D nanostructures consistently exhibiting higher mRNA delivery efficiency than their 2D counterparts. One possible explanation is that planar structures may preferentially adsorb onto the cell membrane, whereas 3D architectures may facilitate membrane wrapping and internalization.

Among the 3D architectures examined, the tetrahedral nanostructure exhibited the highest delivery efficiency. The tetrahedron contains relatively sharp vertices, raising the possibility that local geometric features contribute to membrane interactions and cellular uptake. Sharp nanoscale features could potentially alter local membrane curvature or membrane deformation during carrier-cell interactions. Although further high-resolution fluorescence imaging may be required to directly visualize this relationship, our observations highlight geometric architecture as an important design parameter for mRNA delivery.

In addition to geometry, our platform enables independent incorporation of targeting ligands without altering the underlying nanostructure. Under otherwise comparable conditions, aptamer-functionalized nanostructures exhibited greatly enhanced delivery relative to their non-targeted counterparts, demonstrating that molecular recognition can be integrated with geometric optimization to further improve cellular delivery. This modularity provides an opportunity to combine carrier architecture and receptor-mediated targeting within a single programmable platform.

We further demonstrate that the integration of mRNA into the hybrid nanostructure critically influences delivery performance. Particularly, embedding mRNA within the DNA framework substantially improved delivery compared with attaching mRNA as an external single-stranded cargo. This enhancement may arise from complementary mRNA-DNA duplex formation together with steric confinement within the nanostructure. Previous studies have shown that hybridization with complementary nucleic acids can stabilize mRNA by reducing the exposure of nuclease-sensitive single-stranded regions and protecting sequences from degradation pathways^48, 49^. Our design extends this concept by integrating the hybridized mRNA into a pre-organized nanoscale scaffold. In addition to local duplex protection, the surrounding nanoarchitecture may provide an additional layer of steric shielding that limits accessibility of nucleases to the cargo. Thus, hybridization-mediated protection and nanoscale structural confinement may act cooperatively to improve mRNA stability and functional delivery.

In summary, our results establish a modular design framework in which carrier geometry, molecular targeting, and cargo organization can be systematically tuned to regulate mRNA delivery. This framework is attractive for neoantigen mRNA applications, in which relatively short coding sequences can be integrated with multiple short DNA oligonucleotides to generate a compact carrier-cargo complex-targeting architecture. Importantly, the platform is not restricted to short mRNAs, as we demonstrate the successful delivery of smURFP mRNA, showing its applicability to longer cargo molecules as well. The use of a small number of short DNA oligonucleotides also provides practical advantages in terms of synthesis and assembly. Additionally, crude oligonucleotides could be directly incorporated into the nanostructures after synthesis without intermediate purification based on our study, reducing material loss, processing steps, and associated costs. Overall, this study provides a programmable and experimentally tractable platform for dissecting geometry-dependent mRNA delivery and offers design principles for the development of effective nucleic acid delivery systems.

## Acknowledgements

The authors thank Dr. Hua Wang’s group at UIUC for providing the dendritic cells used in this study. The authors also thank Dr. Adam Friedman and Sherwood Yao at Atom Bioworks Inc. for sharing the proprietary CD33-targeting aptamer sequence used in this study. The authors acknowledge the Imaging Cores at the Materials Research Laboratory (MRL) and the Carl R. Woese Institute for Genomic Biology (IGB) at UIUC for the technical assistance.

